# SKiD: A Structure-Oriented Kinetics Database of Enzyme-Substrate Interactions

**DOI:** 10.1101/2025.05.18.654770

**Authors:** Sowmya Ramaswamy Krishnan, Nishtha Pandey, Rajgopal Srinivasan, Arijit Roy

## Abstract

Enzymes are essential biological catalysts that drive nearly all biochemical reactions. Understanding their efficiency and specificity involves studying enzyme kinetics, particularly the parameters k_cat_ and K_m_. However, there is limited data linking these kinetic parameters with the three-dimensional (3D) structures of enzyme-substrate complexes. Since enzyme function is determined by its structure, such mapping enhances insight into structural basis of enzymatic function and supports applications in enzyme design, synthetic biology and metabolic engineering. To address this critical gap, this work presents SKiD (Structure-oriented Kinetics Database), a comprehensive, structured database integrating k_cat_ and K_m_ values with the corresponding 3D structural data. This is accomplished by integrating data from existing bioinformatics resources using automated programs to process the data and enhancing it with computational predictions. The erroneous data encountered during data integration is manually resolved. Metadata such as literature and assay conditions (e.g., pH and temperature) are preserved. The 3D coordinates of the modelled enzyme-substrate complexes are provided along with their UniProtKB identifier. The database is freely accessible from: https://doi.org/10.5281/zenodo.15355031.

## Background & Summary

Enzymes are biocatalysts that aid biochemical reactions by reducing the activation energy barrier of substrate to product conversion and increasing the rate of reaction. Thus, enzymes are crucial modulators of important biological processes like cellular metabolism and cell signalling. A system-level study of complex biological networks requires detailed kinetic information about each enzyme-catalysed biochemical reaction (Zhou et al., 2021). The rate of enzymatic reaction is expressed as a function of the substrate concentration with the help of two constants, turnover number (k_cat_) and the Michaelis-Menten constant (K_m_) (Michaelis et al., 2011). These parameters are fundamental for understanding enzyme kinetics and evaluating their catalytic efficiency (Koshland, 2002). Driven by environmental sustainability concerns, there is a growing interest in the study of enzymes for potential applications in the chemical and pharma industries (Savile et al., 2010; Sheldon and Woodley, 2018). Computational models of enzyme-catalysed reactions are important tools for industries that use biocatalysts (Česnik et al., 2020). Hence, multiple databases that store enzyme kinetic parameters have been developed to ensure that application of biocatalysts can be successful in engineering better biocatalysts with higher efficiency and selectivity.

SABIO-RK database was developed in 2006 to provide high quality enzyme kinetics data to researchers. The data is extracted from literature through manual curation, prioritizing quality over quantity (Wittig et al., 2018). BRENDA (BRaunschweig ENzyme DAtabase) on the other hand serves as the most comprehensive resource for enzymes (Placzek et al., 2017). The enzyme-substrate kinetic data in BRENDA, is retrieved from scientific literature using KENDA’s (Kinetic ENzyme Data) automated text mining approach. The BRENDA 2016 database version had ∼8,500 kinetic values mined from 11,886 literature (Placzek et al., 2017). However, despite recommendations from the Standards for reporting enzymology data (STRENDA) commission, enzyme kinetic data is not always documented unambiguously in the literature (Tipton et al., 2014), leading to metadata annotation issues in automated text mining. The development of STRENDA DB in recent times is an attempt to ensure appropriate reporting of kinetic data by the researchers themselves (Swainston et al., 2018). However, not every biochemistry journal requires an SRN (STRENDA Registry Number), so data submission to STRENDA DB is at the discretion of researchers and the data archived is incomplete. ProtaBank is a new information resource proposed as a central repository for protein engineering data (Wang et al., 2018). It contains information from 1671 studies, but as the name suggests, it is not solely dedicated to enzymes or kinetic data. Further, the information might be experimental, computational or derived. GotEnzymes (Li et al., 2023), another database with predicted values, stores millions of enzyme kinetic data points generated using the deep learning-based method *DLKcat* (Li et al., 2022). However, the performance of *DLKcat* method has been critically evaluated and concerns have been raised on the quality of the predicted data (Kroll and Lercher, 2024). Hence, a curated kinetic database derived from BRENDA, the most comprehensive resource, can be considered as an important contribution. Still, the challenge is to map the kinetic data to the corresponding three-dimensional structure of the enzyme-substrate pair which can provide details of enzyme-substrate intermolecular interactions.

Literature evidence suggests that enzyme activity and kinetic properties show better correlation with their three-dimensional structure compared to sequences (Qian et al., 2025). For example, structure of the serine protease significantly influences its kinetic parameters. The precise spatial arrangements of amino acids in the binding site, specifically that of the catalytic triad (Ser, His, Asp) determines the enzyme’s substrate specificity and catalytic efficiency (Perona and Craik, 1997; Borkakoti et al., 2025). Similarly, phylogenetic analysis of the *E. coli* haloacid dehalogenase-like hydrolases superfamily members using sequences versus catalytic efficiency showed lack of congruence (Kuznetsova et al., 2006). However, structure analysis showed presence of common scaffolds conserved among the family members (Burroughs et al., 2006). Therefore, integrating structural information with substrate-specific enzyme kinetic data would be a value addition towards understanding how the enzyme interacts with substrates and correlates with catalytic efficiency. This information, presented in the form of enzyme substrate complex structure, can serve as a useful resource in designing improved enzymes for industrial purposes. IntEnzyDB (Yan et al., 2022) is a resource where both the enzyme structure and kinetic information were curated. However, it contains only 1050 enzyme-substrate pairs, where kinetic data is mapped to 155 unique protein structures. It provides substrate details and enzyme PDB ID as separate columns, along with active site residues. This information is extracted from the UniProtKB annotations, which are often inferred from PROSITE-ProRule annotation or sequence similarity (Sigrist et al., 2005).

In this work, we have developed a Structure-oriented Kinetic Database (SKiD) that reposits kinetic parameters of enzyme-substrate pairs along with their three-dimensional structures by integrating data from several resources. The dataset generation involved the use of multiple computational methods. We have reported the challenges faced in the process and described how they were overcome. Although, we focused on automating the process as far as possible, manual intervention was required at multiple steps. SKiD consists of 13,654 unique enzyme-substrate complexes spanning six enzyme classes. The dataset includes the activity of both wild-type and mutant enzymes measured with natural as well as non-natural substrates. The datapoints have been generated after extensive pre-processing e.g., protonation based on experimental pH, and is ready for use in several downstream applications. This is the first and only such resource to the best of our knowledge. The database is freely accessible from: https://doi.org/10.5281/zenodo.15355031.

## Methods

The Michaelis-Menten constant (K_m_) and enzyme turnover number (k_cat_) were considered as the primary measures reflecting enzyme-substrate interaction kinetics for this study. To construct the database, the K_m_ and k_cat_ values of wild-type and mutant enzyme-substrate interactions were curated from the BRENDA database (Chang et al., 2021). The PDB structures of enzymes were mapped based on BRENDA-derived UniProtKB annotations, wherever available. The substrate and cofactor binding sites were mapped for each enzyme based on the bound ligand or its functional homologue. Mutant enzymes were modelled from their wild-type structures and the protonation states of all enzymes were corrected based on the experimental pH and temperature from BRENDA.

The pre-processed enzyme and substrate structures were docked to obtain the final database of the enzyme-substrate complex structures. The complete workflow followed for development of the database is shown below (Fig. 1).

**Figure 1:**
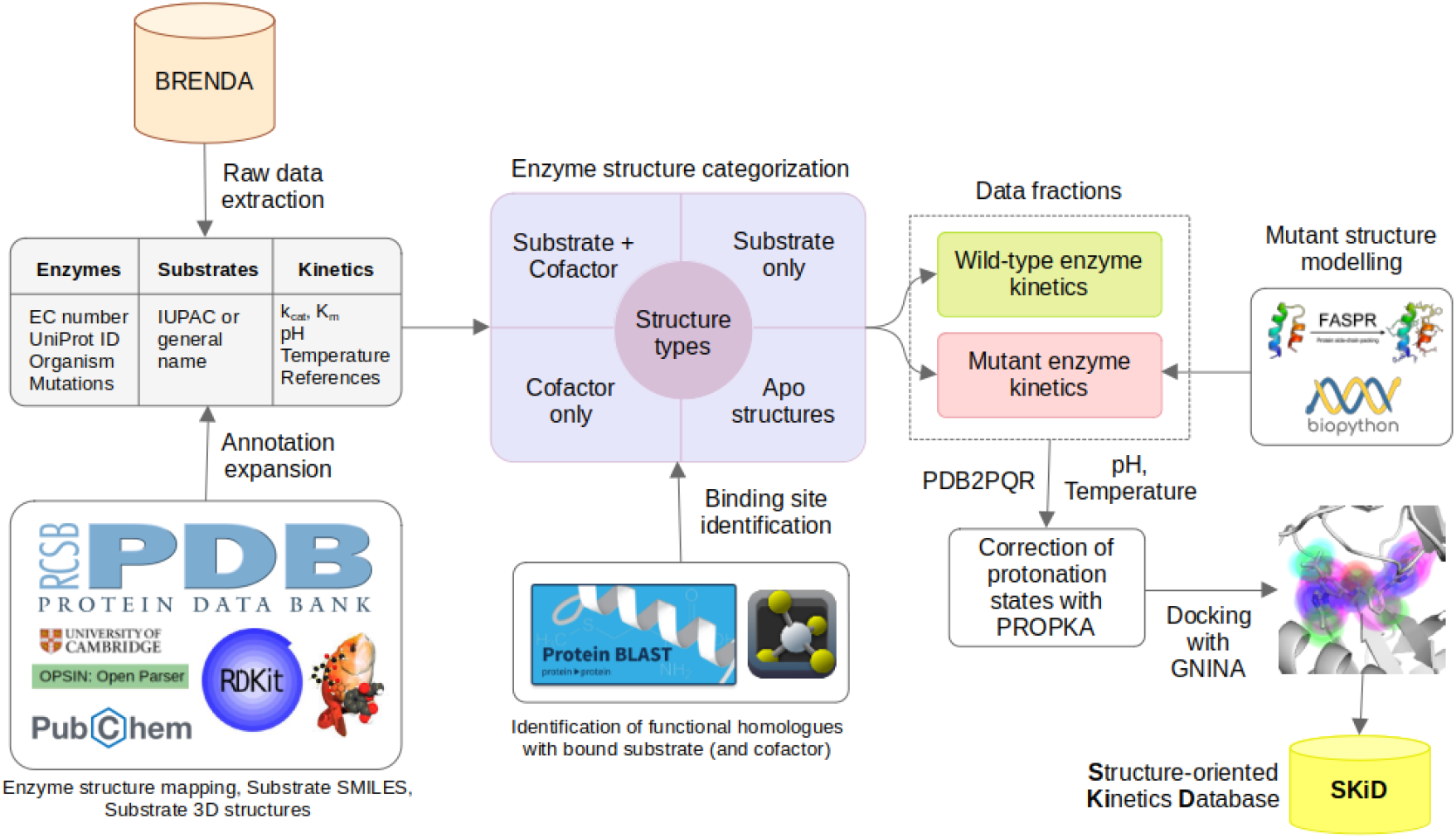
Complete workflow followed for the development of SKiD. Each step of data processing and the external databases and tools utilized for the workflow are also provided.

### Enzyme-substrate interaction kinetics data curation

Data on experimentally measured K_m_ and k_cat_ values of enzyme-substrate pairs were collected from the BRENDA (v2023) database (Chang et al., 2021). In-house scripts were used to process the raw data from the database and arrive at a uniform format for database development. Redundancy in the dataset was resolved through extensive comparisons of the annotations extracted for each datapoint including, enzyme commission (EC) number, UniProtKB ID of the enzyme, substrate SMILES, wild-type or mutant form of the enzyme for which the kinetics data was measured, experimental conditions such as pH and temperature, and the cited reference for the K_m_ and/or k_cat_ values. When multiple datapoints for the same enzyme-substrate complex under same experimental conditions were found to have differing K_m_ and k_cat_ values, their geometric mean was reported in the database in accordance with previous studies (Heckmann et al., 2018; Kroll et al., 2023; Yu et al., 2023). In case of redundant datapoints with a wide range of values (minimum value was lower than half of the maximum value), the geometric mean was computed post manual verification of the values from supporting literature. An outlier analysis was also performed on the K_m_ and k_cat_ values to prune datapoints with values outside thrice the standard deviation of the log-transformed parameter distributions. These datapoints are provided in the Supplementary Information 2.

### Enzyme and substrate annotations

Next, enzyme and substrate annotations were performed. This is an important step to identify their 3-dimensional structure. BRENDA can contain multiple annotations specific to the enzyme and substrate. Datapoints were also found to have two different annotation formats namely, comments and named columns (Supplementary Information 1 - Fig. S1). For enzymes, the annotations were extracted from comments and other named entries in the database through custom Python scripts. Experimental conditions and mutations in the enzyme were only available in the comments section from which, regular expressions were used to extract and standardize this data. The PDB IDs of the enzymes were extracted from their UniProtKB IDs to map the enzyme structure to each datapoint. The IUPAC names of substrates in BRENDA were extracted and annotated with their isomeric SMILES using the OPSIN (Lowe et al., 2011) and PubChemPy libraries. Non-small molecular substrates such as proteins, peptides, nucleic acids, polymers and peptide-sugar conjugates were omitted from the dataset. Since several substrates had non-standard nomenclature in BRENDA, extensive manual annotation from several standard databases including PubChem (Kim et al., 2023), ChEMBL (Zdrazil et al., 2024), ZINC (Tingle et al., 2023), and ChEBI (Hastings et al., 2013) was also undertaken to increase substrate coverage of the database. Additionally, for the substrates that remained unresolved from the previous step, SMILES were generated by drawing structures using the GChemPaint tool (Bréfort, 2001) based on the product catalogue of laboratory suppliers like Thermo Fisher Scientific. Finally, the three-dimensional structure of substrates was obtained from their SMILES using the RDKit (https://www.rdkit.org) and OpenBabel (O’Boyle et al., 2011) packages. Explicit hydrogens were added, and the structures were subjected to a short energy minimization using the MMFF94 force field (Halgren, 1996).

### Mapping the structure of known enzyme-ligand complexes to the kinetics dataset

The crystallographic structures of the enzyme-substrate pairs might not be available for majority of the kinetic parameters collected in the previous step. Various strategies were adopted to obtain these structures. Initially, the available structural information was collected by mapping the PDB structures based on UniProtKB annotations for each enzyme in the dataset. These structures were classified into four categories: substrate+cofactor structures, substrate-only structures, cofactor-only structures and apo structures since they need to be treated differently for the mapping. Crystal additives and ions present in the structures were not considered as substrates or cofactors during the analysis. The substrates were differentiated from cofactors in each PDB structure using the mapping available from the EMBL CoFactor database (Fischer et al., 2010). After the automated discrimination of substrates from cofactors, manual verification was performed and cases where cofactors can act as substrates were identified and corrected. Several entries in CoFactor database were also found to be incomplete, leading to wrong categorization of the structures. Such cases were corrected manually using the literature associated with the structure, wherever available.

Once the PDB structures were categorized, each datapoint in the database was associated to multiple structures from different categories. In such cases, datapoints with at least one substrate+cofactor or substrate-only structure mapped from PDB were considered as successful entries. Few hundreds of entries were however associated to only apo structures or cofactor-only structures. In such cases, the structures were mapped to the closest functional homologues with bound substrate using BLASTp (Altschul et al., 1990) against the PDB database with an e-value cut-off of 0.01. If such close homologues could not be identified, distant functional homologues with bound substrate were mapped through PDB advanced search and manual structural alignment using backbone RMSD cut-off of 3 Å. Only those cases were considered where at least 90% of the binding site residues remain conserved between the parent structure and the homologue. In case of enzymes with multiple known PDB structures, preference was assigned based on resolution and completeness of the structure. The structure mapping obtained for all the enzymes present in BRENDA database along with their categories are available in the Supplementary Information 2.

### Extraction of cognate substrate and cofactor binding sites from mapped structures

In accordance with the sc-PDB (Desaphy et al., 2015) convention, the substrate and cofactor binding sites were defined as residues with at least one atom within 6.5 Å from the substrate and cofactor atoms, respectively. For enzymes mapped to at least one substrate+cofactor or substrate-only structures, the binding sites were directly identified from the crystal structure using BioPython (Cock et al., 2009). For enzymes that had cofactor-only or apo structures with close or distant functional homologues, the bound substrate from the homologous structure was utilized to extract the binding sites. This was achieved by alignment of the homologous structure to the parent structure using the PyMOL API from Python, and saving the coordinates of the substrate post-alignment with the coordinates of the parent structure. In cofactor-only structures, the cofactor from the parent structure and the substrate from the functional homologue were used to map the binding sites. In apo structures, the binding sites were mapped depending on the ligands bound to the functional homologue chosen from PDB.

### Mapping the mutations reported in BRENDA to the PDB structures

The dataset collected from BRENDA includes both wild-type and mutant enzymes. However, the position of the mutated residue reported in BRENDA is predominantly based on the UniProtKB sequence position, which does not necessarily coincide with the residue numbering followed in PDB structures. Hence, the EMBL SIFTS database (Dana et al., 2019) was used to derive the residue position maps between UniProtKB sequences and PDB structures. Based on the unresolved chain-level SIFTS numbering maps, 15 error types were identified during mutant position mapping in enzyme structures (Table 1) and resolved using custom scripts. The distribution of these error types across BRENDA is provided below (Fig. 2). Few of these error types and their corresponding examples from BRENDA are discussed in the Results section. The error type mapping for all the mutant enzymes reported in BRENDA database is also provided in the Supplementary Information 2. After mapping the mutated residue positions from BRENDA to PDB structures, mutations were further classified into binding site and non-binding site mutations based on the binding site definition discussed earlier.

**Table 1:** Errors encountered during mapping of mutated residue positions from BRENDA dataset to enzyme structures from PDB. For each error type, an error code and the solution type followed are also provided.

| Error code | Error type | Solution type |
| --- | --- | --- |
| 1 | Wrong UniProtKB ID linked in BRENDA | Automated |
| 2 | Wrong mutation data curation in BRENDA | Manual |
| 3 | UniProtKB ID mismatch between BRENDA and PDB | Automated |
| 4 | Isoforms and INS/DEL variants in BRENDA comments | Manual |
| 5 | No UniProtKB ID linked to PDB (or) no SIFTS mapping available | Automated |
| 6 | Typo in mutation – regex issues in mutation extraction | Automated |
| 7 | Sequence mismatch between UniProtKB and BRENDA/Reference article | Manual |
| 8 | Reverse alignment issue – UniProtKB sequence offset from PDB sequence | Automated |
| 9 | Residue numbering in non-standard fashion | Manual |
| 10 | Mutated PDB structure – WT residue not matching to the UniProtKB in reverse alignment | Automated |
| 11 | Partial PDB structure – mutated residue not part of resolved structure | Automated |
| 12 | Residue numbered in sequential order – not based on author-provided residue numbering | Automated |
| 13 | Modified residue (non-standard residue or L-amino acid) used for mutation | Automated |
| 14 | Continuous residue numbering between PDB chains in multimeric enzymes | Manual |
| 15 | Unexplained | Automated |

**Figure 2:**
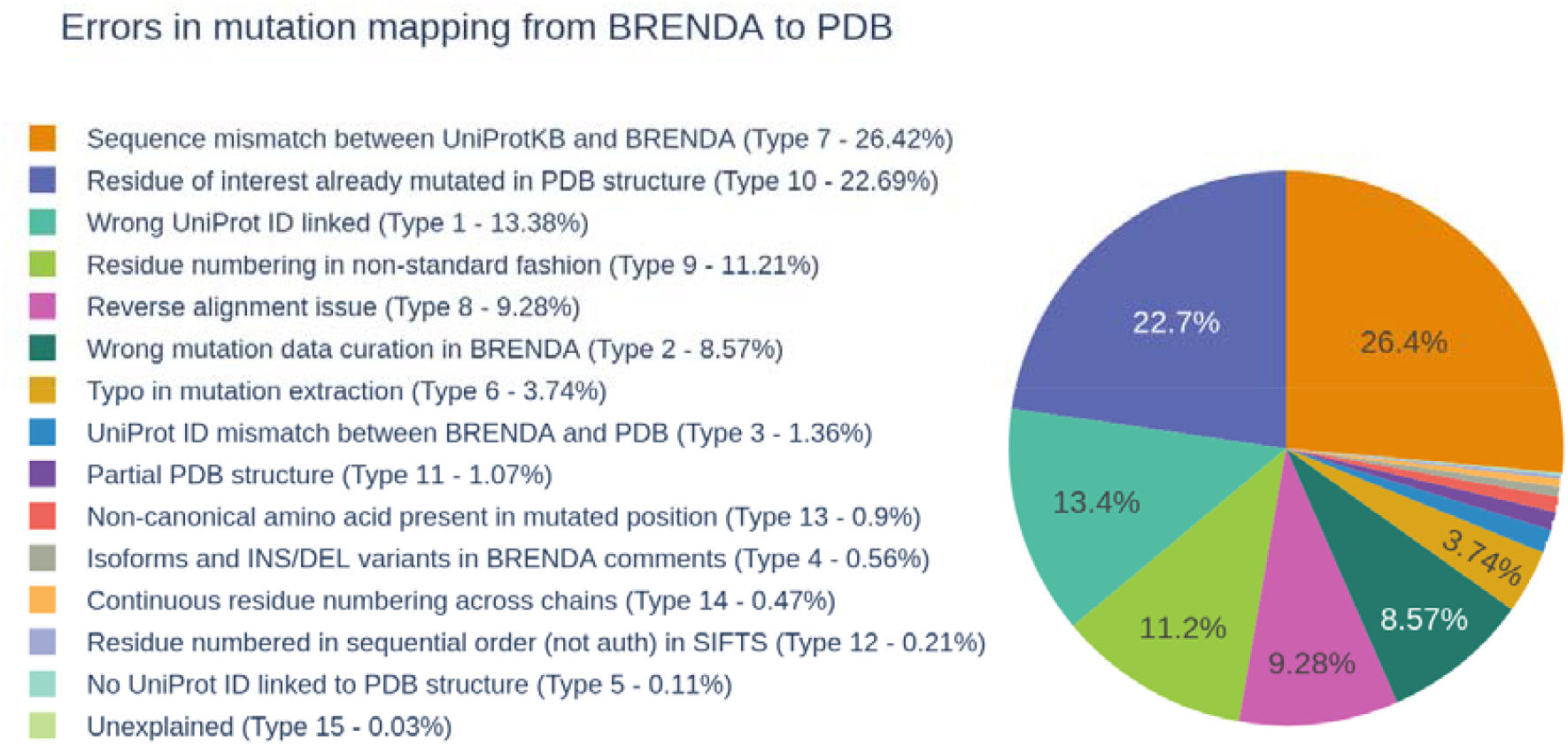
Percentage of occurrence of the error types identified in BRENDA database during mutation mapping from BRENDA to PDB structures.

### Modelling mutant structures with FASPR program

While several mutant enzyme structures mapped from PDB had the exact mutation reported in the BRENDA database, all the mutant structures were modelled from their wild-type structures to maintain the dataset consistency. Two computational tools for mutant structure generation from the experimental wild-type structure – FASPR (Huang et al., 2020) and SCWRL4 (Krivov et al., 2010), were tested on a set of 10 experimentally resolved mutant enzyme structures. The wild type PDB structures were used as the starting conformation and mutant conformations were generated using either position specific side chain repacking or all positions side chain repacking. The performance of FASPR and SCWRL4 was compared using all atoms RMSD value of position specific in silico mutated structures with their experimentally resolved structures as reference. ProFit server was used to compute all atoms RMSD values (McLachlan, 1982), the fit was centred around mutant site; owing to the higher flexibility of terminal residues, 10 terminal residues from both sides of the polypeptide chain were uniformly excluded from the calculations. Based on the comparison and another existing benchmark study (McPartlon and Xu, 2023), FASPR was used for mutant structure generation. It was also used to replace missing atoms and residues using the OpenMM package (Eastman et al., 2024) in the background for repairing the enzyme structures. FASPR can perform multiple mutations at once in the same structure, which was effectively leveraged to model the mutant enzymes reported in BRENDA database. For enzymes without wild-type structures in PDB, the available mutant structure was reverted to the wild-type structure by systematically mutating using FASPR with the UniProtKB-derived sequence as the wild-type reference.

### Modification of the protonation state of enzyme structures based on experimental data from BRENDA

The experimental data included for each enzyme-substrate interaction in BRENDA includes the organism, pH and temperature of interest. While the UniProtKB ID is unique to an enzyme-organism pair, the pH effect was also incorporated into the enzyme structure through the protonation state. The PROPKA program (Olsson et al., 2011) available as part of the PDB2PQR suite (Dolinsky et al., 2007) was used to modify the protonation state of polar residues in the wild-type and mutant enzyme structures. Manual literature curation was performed to retrieve pH information if the information was missing and for the remaining cases pH 7 was used to determine the default protonation state. Since chemical reactions involved in the biosynthesis of organic molecules is highly dependent on proton addition and removal, this pre-processing step is invaluable for atomistic computational studies such as docking and molecular dynamics simulations on the enzymes.

### Modelling the enzyme-substrate interactions through docking

With the pre-processed enzyme and substrate structures, the enzyme-substrate complexes were modelled through molecular docking. If the enzyme was mapped to at least one substrate+cofactor complex in PDB, the cofactor was retained during the docking calculations to effectively constrain the docking grid covering the substrate binding site so that the possible substrate-cofactor interactions can be captured. To perform the docking calculations, the GNINA (McNutt et al., 2021) and SMINA (Koes et al., 2013) programs were initially benchmarked using the Platinum database (Pires et al., 2015), which contains experimentally determined wild-type and mutant protein-ligand complexes along with their binding affinity values. Two different experiments were performed wherein either a single docking pose or multiple docking poses were sampled by the programs. The similarity between the docking pose(s) and the ligand pose from the crystal structure were compared using two measures: RMSD and ligand centroid distance. While RMSD captures the overall agreement between the poses, the ligand centroid distance can capture the position of the docking grid accurately, even if the orientation of the ligand is different due to the numerous rotatable bonds present in flexible ligands. More details on the benchmarking are provided in the Results section. Based on the benchmarking, GNINA program with multiple pose sampling and CNN re-scoring was chosen. The docking grids were set using the *autobox_ligand* option, which uses the already bound substrate as the reference ligand for grid setting. The top-ranked docking pose out of 10 generated poses was utilized for generation of the final enzyme-substrate complex structures.

## Data Records

The SKiD database is freely accessible as a collection of spreadsheets and a structure archive via Zenodo: https://doi.org/10.5281/zenodo.15355031. The database includes two spreadsheets containing the k_cat_ and K_m_ datasets separately, since BRENDA contains hundreds of instances with either only the k_cat_ value or only the K_m_ value. The unique enzymes and substrates present in the database are also provided with appropriate annotations as additional spreadsheets. Each datapoint in the database is described using 16 columns (Table 2), preserving the annotations available in the BRENDA JSON file. Depending on the application of interest, users can filter subsets of data from the spreadsheets along with their corresponding structure models from the archive. Each entry in SKiD is provided a separate entry number, which is repurposed as a folder label in the structure archive for ease of use. Each folder in the structure archive includes the FASPR-optimized enzyme structure protonated as per the experimental pH value, the substrate docking pose in SDF format, and a meta-data file including details of the organism, substrate SMILES, UniProtKB ID, kinetic parameters etc. The file name in each folder includes the PDB ID, pH value and molecule ID to facilitate comparison of same substrate binding to multiple enzymes and enzyme-substrate complexes tested under different pH conditions. A separate archive of numbered references is also provided for each EC number to preserve the bibliography system used in BRENDA.

**Table 2:** Columns describing each datapoint in the SKiD database with a brief description of the contents within each column.

| Column name | Description |
| --- | --- |
| Entry_ID | A unique identifier provided to each data point in the SKiD database |
| EC_number | Enzyme Commission (EC) number of the enzyme |
| Substrate | Generic or IUPAC name of the substrate. The substrate name provided in BRENDA is preserved to map back to the reaction easily. |
| UniProt_ID | UniProtKB identifier of the enzyme |
| Protein_file | Name of the enzyme structure file in the structure archive of the SKiD database |
| Organism_name | Name of the parent organism (enzyme source) |
| kcat_value (or)<br>Km_value | Value of the kinetic parameter of interest ($k_{cat}$ or $K_m$) |
| Mutant | A binary identifier to discriminate mutant enzymes from wild-type enzymes |
| Mutation | The mutation(s) present in the enzyme as reported in the BRENDA database. Multiple mutations are separated by a forward slash (/) symbol. |
| pH | pH of the reaction as reported in BRENDA database or supporting literature |
| Temperature | Temperature of the reaction (in °C) as reported in BRENDA database or supporting literature |
| References | A list of supporting literature as reported in BRENDA database |
| Site_type | One of 4 types of active site in SKiD database (Substrate+Cofactor, Cofactor only, Substrate only, Apo structure) |
| Substrate_SMILES | Isomeric SMILES of the substrate |
| Mol_file | Molecule identifier and name of the substrate file in the structure archive of the SKiD database |
| pkcat_value (or)<br>pKm_value | Log-scale ($\log_{10}$) value of the kinetic parameter of interest |

## Technical Validation

### Database statistics and feature coverage analysis

The SKiD database includes the structures and kinetic parameters of 13,654 unique enzyme-substrate complexes, derived from 8,562 unique enzymes and 3,248 unique small molecular substrates. There are enzymes from 688 unique organisms in the database. The numbers of unique wild-type and mutant enzymes are 2,169 (25.33%) and 6,393 (74.66%), respectively. The number of unique binding site mutations observed were 2,356 and non-binding site mutations observed were 2,401. Several enzymes were found to include both types of mutations in their structure and a maximum of 12 mutations tested in parallel (Supplementary Information 1 - Fig. S2).

The range of log-transformed k_cat_ (pk_cat_) and K_m_ (pK_m_) values covered by the database is shown in Fig. 3. In terms of the enzyme diversity, the first six classes of enzymes are covered by the database (Fig 3c), with class 1 (oxidoreductases) being the most-represented class and class 6 (ligases) being the least-represented class. The internal diversity metric computed for the substrate dataset using ECFP4 fingerprints was found to be 0.64, indicating high substrate diversity. To visualize the substrate diversity, a TMAP was constructed using the ECFP4 fingerprints derived from substrate SMILES (Supplementary Information 1 - Fig. S3). While there are strong substrate clusters in local branches across the TMAP, especially for EC classes 1 and 3, global clustering of substrates was not visible, which also reinforces the high substrate diversity in the database. An analysis of the effect of mutations on enzyme kinetic parameters is also included in the Supplementary Information 1 – Section S1.

**Figure 3:**
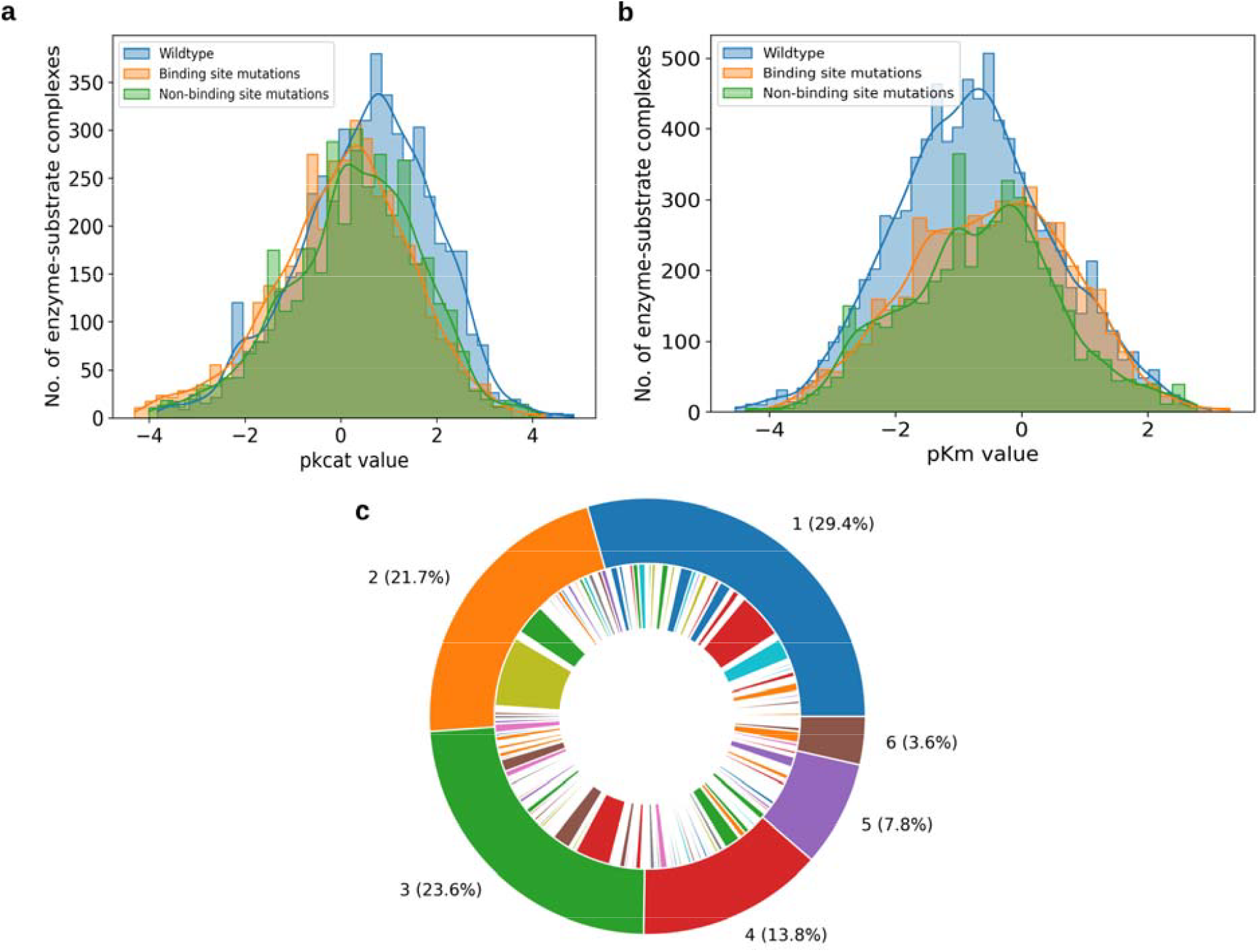
Distributions of (**a**) pk_cat_, (**b**) pK_m_, and (**c**) EC classes in the SKiD database. The outer ring of the nested pie chart shows the total number of datapoints under each EC class while, the inner ring shows the number of unique organisms covered by the EC class.

### Annotation of substrate binding site requires manual intervention

As mentioned in the Methods section, the PDB structures mapped to the enzyme-substrate complexes curated from BRENDA were categorized into 4 types: substrate+cofactor structures, substrate-only structures, cofactor-only structures and apo structures (Table 3). Based on Table 3, majority of the structures belonged to the substrate-only category in both the datasets. The annotation of substrate binding sites for substrate+cofactor and substrate-only structures was done directly by extraction of residues lying within 6.5 Å radius from the substrate and/or cofactor. In case of structures with multiple bound ligands (as shown in Supplementary Information 1 – Fig. S5), the literature associated with the PDB structure was manually curated to identify the ligand bound at the active site (substrate). Except the substrate and the cofactor, any other ligand bound to the enzyme was omitted from further analysis. For cofactor-only and apo structures, homologous enzymes were initially identified through BLASTp search for annotation of the binding site.

**Table 3:** Statistics of the four categories of PDB structures in the k_cat_ and K_m_ datasets curated from BRENDA. The numbers are based on only the wild-type structures in the dataset.

| Structure category | $k_{cat}$ dataset | $K_m$ dataset |
| --- | --- | --- |
| Substrate + Cofactor | 426 | 543 |
| Substrate-only | 1155 | 1442 |
| Cofactor-only | 29 | 42 |
| Apo structures | 11 | 86 |

Upon mapping the substrate binding sites from homologues for cofactor-only structures, both proximal and distal sites from the cofactor could be observed. Distal substrate binding sites (substrate lies outside the 6.5 Å radius) were observed either due to distinct domains (Fig. S6a) for the substrate and cofactor or due to tunnel-shaped binding sites in the structure (Fig. S6b). In case of aminoaldehyde dehydrogenase (EC 1.2.1.19) from *Pisum sativum* (PDB ID: 3IWJ), the binding site mapping from the homologous structure (PDB ID: 4I9B) led to the identification of a tunnel-shaped binding site with the substrate and cofactor bound to extreme ends of the tunnel (Fig. S6b). Since distal substrate binding sites can be due to crystallization artifacts (Pochapsky and Pochapsky, 2019) or exit state of the reaction product (Kingsley and Lill, 2015), such cases were not included in the SKiD database and were marked as unresolved.

On the other hand, proximal substrate binding sites in cofactor-only structures also included certain special cases. Covalent substrate-cofactor interaction was observed in the case of phosphoserine aminotransferase (EC 2.6.1.52) from *Niallia circulans* (PDB ID: 2C0R) upon mapping to its structural homologue (PDB ID: 4AZJ) (Fig. S7a). Significant movement in the cofactor head group position could also be observed upon substrate binding as in the case of Ferridoxin-NADP+ reductase (EC 1.18.1.2) from *Xanthomonas citri* (PDB ID: 4B4D) (Fig. S7b). These proximal substrate binding sites involving covalent bond formation and significant cofactor movement were also omitted from further analysis. While these observations indicate the magnitude of variation in substrate processing across homologous enzymes in the SKiD database, they also indicate the difficulty in reliable automation of binding site mapping across enzymes through structure homology. This necessitates manual processing and careful verification of the mapped substrate binding sites prior to further structural studies, as undertaken in this database.

### Challenges in mutation mapping from BRENDA to PDB structures

While enzymes in BRENDA are mapped to their corresponding UniProtKB IDs, multiple errors (Table 1) and challenges were encountered, which resulted in residue numbering mismatch between UniProtKB sequence and PDB structures. This hampered direct mutation mapping from BRENDA to PDB. Representative examples of the challenges encountered during mutation mapping are tabulated below (Table 4). These examples highlight the hurdles encountered in automated mutation mapping between experimental studies as documented in BRENDA database with enzyme structures from PDB. SKiD aims to provide highly reliable mutation mapping for structure-based enzyme kinetics studies through manual verification of every unresolved datapoint in BRENDA.

**Table 4:** Representative examples of various errors encountered during mutation mapping from BRENDA database to the enzyme PDB structures. The error codes as listed in Table 1 are provided for each example, along with a brief description of the error.

| EC number | Enzyme | Organism | Mutation(s) reported in BRENDA | Error/Challenge encountered | Error code |
| --- | --- | --- | --- | --- | --- |
| 1.1.1.203 | Uronate dehydrogenase | *Chromohalobacter salexigens* | D236N | Wrong mutation mapping in BRENDA – should be D246N according to the cited literature | 2 |
| 1.1.1.27 | L-lactate dehydrogenase | *Geobacter stearothermophilus* | D38SC81S | Typo in mutation reported – should be D38S/C81S | 2 |
| 1.1.1.205 | IMP dehydrogenase | *Mycobacterium tuberculosis* | MtbIMPDH2 DELTACBS | Non-standard format of mutation in BRENDA | 4 |
| 1.1.1.205 | IMP dehydrogenase | *Homo sapiens* | A285T | UniProtKB ID mismatch between BRENDA (P20839) and PDB (Q5H9Q6) resulting in isoform difference | 5 |
| 1.1.1.267 | 1-deoxy-D-xylulose-5-phosphate reductoisomerase | *Mycobacterium tuberculosis* | W203Y | UniProtKB ID is linked to BRENDA (P9WNS1), but not linked to PDB ID (2Y1F), resulting in absence of SIFTS mapping for the structure | 5 |
| 1.1.1.363 | Glucose-6-phosphate dehydrogenase | *Leuconostoc mesenteroides* | K21R | The mutation can be directly traced back to PDB structure (PDB ID: 1DPG), but not to UniProtKB sequence (P11411) due to a missing Methionine residue in N-terminal | 8 |
| 1.1.1.37 | Malate dehydrogenase | *Haloarcula marismortui* | R100Q | Mutation numbered in reference study based on the position from a multiple sequence alignment | 9 |
| 1.1.1.119 | Glucose-1-dehydrogenase | *Sulfurisphaera tokodaii* | V54I, I189V | Only partial structures available in PDB | 11 |
| 1.11.1.19 | Dye decolorizing peroxidase | *Irpex lacteus* | W109F | Mutation does not follow either UniProtKB sequence (A0A2P1C6N4) or PDB (7D8M) residue numbering. Instead follows sequential residue numbering in SIFTS | 12 |

### Results from benchmarking of mutated structure modelling tools

The structures of engineered enzymes were not available for all datapoints and hence, the mutant conformations were predicted *in silico*. The FASPR method for side chain repacking performed reasonably well as quantified by the deviations between modelled and solved structures (Table S3). Similar findings have also been reported by a previous study (McPartlon and Xu, 2023). FASPR not only performed at par with other recently developed methods, but also, emerged as the fastest tool. These results made FASPR the natural choice for processing large datasets such as BRENDA. The scope of repacking in a position specific manner versus for all residues was also explored while generating the mutant conformations. Repacking all positions led to larger RMSD values and significant conformational changes within an 8 Å radius around the mutation site when compared with the reference structure. Hence, in the subsequent steps, only position specific repacking was executed on the larger dataset. An example comparing the effect of different repacking modes available in FASPR is provided in Supplementary Information 1 (Fig. S8).

### Results from benchmarking of docking programs for modelling enzyme-substrate complexes

Since the experimental structures of more than 90% of the enzyme-substrate complexes in SKiD were not available from PDB, they were modelled using molecular docking calculations. With the availability of several docking programs for protein-ligand complexes, selection of the suitable docking program and the best docking pose are non-trivial problems (McNutt and Koes, 2015). Hence, a benchmarking was performed using two state-of-the art programs (GNINA and SMINA) on the Platinum database (Pires et al., 2015). GNINA (McNutt et al., 2021) offers CNN-based re-scoring of docking poses along with the traditional biophysical scoring function and the ability to accelerate pose generation with GPUs. On the other hand, SMINA (Koes et al., 2013) uses only the traditional scoring function with multi-core CPUs to achieve similarly accurate binding poses.

The Platinum database is composed of 1,008 non-redundant wildtype-mutant protein pairs bound to the same ligand with change in their ligand binding affinity known from experiments. However, only 169 structure pairs were mapped to their respective PDB structures in the database and these were used as the benchmarking dataset for the docking programs (Supplementary Information 2). Prior to docking, the ligand conformation and coordinates were randomized using RDKit to mimic the real-world docking scenario. Based on a comparison of the RMSD of the docking poses to the experimental ligand conformation for the wild-type proteins, GNINA was observed to have first docking pose as the best pose in 67.46% of the calculations (Fig. 4a). In comparison, SMINA had the first docking pose as the best pose in only 44.57% cases. In terms of the difference in RMSD with the experimental ligand conformation between the first and best docking poses (Fig. 4b), GNINA had a lower average difference (0.71 Å) compared to SMINA (2.16 Å) indicating that the first docking pose from GNINA (with CNN rescoring) can be used as a reliable proxy for the best docking pose during enzyme-substrate docking calculations.

**Figure 4:**
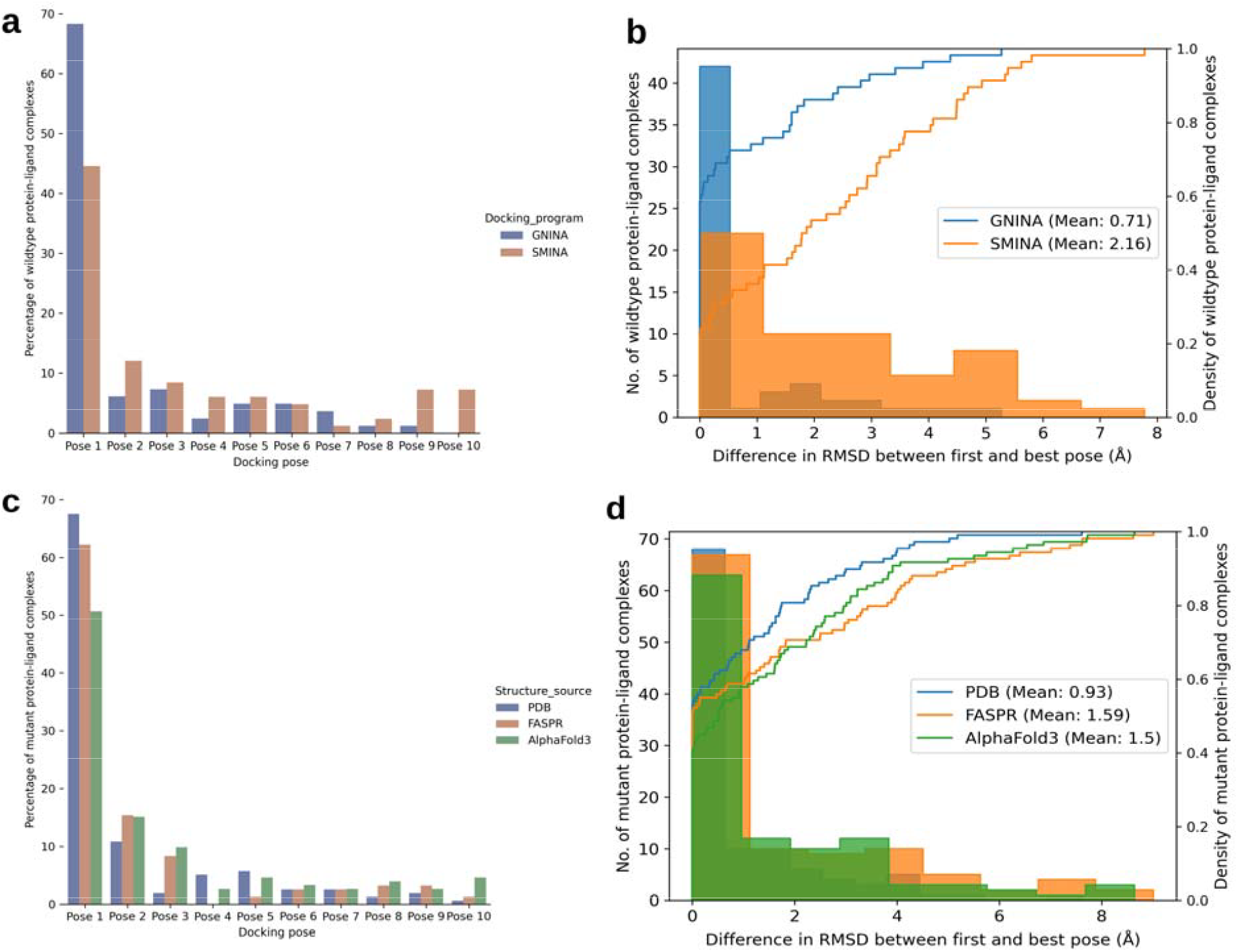
Comparison of docking results between GNINA and SMINA with the Platinum database (**a**) Percentage of wildtype protein-ligand complexes for which each of the top 10 sampled poses is the best pose in terms of RMSD; (**b**) Difference in RMSD between first and best docking poses upon comparison with the crystal pose. Comparison of docking results for different sources of mutant structures (PDB, FASPR and AlphaFold3) with the Platinum database (**c**) Percentage of mutant protein-ligand complexes for which each of the top 10 sampled poses is the best pose in terms of RMSD; (**d**) Difference in RMSD between first and best docking poses upon comparison with the crystal pose.

Since the structures of mutant proteins in the database are not from experiments, the benchmarking was also extended to modelled mutant structures from FASPR (Huang et al., 2020) and AlphaFold3 (Abramson et al., 2024) using GNINA. Specifically, the conformations from AlphaFold3 have been previously shown to be in between the true apo and holo states of the enzyme (Yasumitsu and Ohue, 2025), increasing the difficulty in rigid docking calculations. Comparison of the RMSD of the 10 docking poses with the crystal pose between experimental and modelled mutant structures (Fig. 4c) indicates that with FASPR-derived mutants, GNINA-derived first docking pose is the best docking pose in a large fraction of the dataset (62.17%). It also remains comparable to the results from experimental mutants wherein, the first docking pose is the best pose in 67.51% cases. But for AlphaFold3-derived mutants, this fraction is only 50.65%, although the average difference in RMSD with the crystal conformation between the first and best poses is lower than that from FASPR-derived mutant structures (Fig. 4d). Hence, utilizing FASPR-derived mutant structures as input for enzyme-substrate docking was found to be closer to using the experimental mutant structures due to the ease of docking pose selection, compared to AlphaFold3. From the comparison, it can also be inferred that local structural changes made by FASPR do not significantly impact the holo conformation in comparison with *ab initio* structure prediction methods such as AlphaFold3.

Based on these results, GNINA with multi-pose sampling and CNN rescoring was found to outperform SMINA with both experimental and modelled enzyme structures. The results from GNINA and SMINA for single pose sampling with and without rescoring are provided in the Supplementary Information 1 (Fig. S9) for the Platinum database. An example of high and low RMSD docking poses from GNINA and SMINA is also provided in Supplementary Information 1 (Fig. S10).

### Potential applications for the SKiD database

Since the SKiD database has a uniquely curated dataset of enzyme-substrate complex structures and their associated enzyme kinetics parameters from experiments, it will be an invaluable resource for the enzyme engineering community. Several studies have attempted to engineer natural enzymes for green chemistry (Radley et al., 2023; Paul et al., 2024), to aid large-scale synthesis of drugs and industrially useful molecules through bacterial systems (Sheldon et al., 2020; Siedentop and Rosenthal, 2022). Mutations favoring improvement in kinetic properties of enzymes from the database can be used to design targeted mutagenesis studies focused on enzyme activity optimization. On the other hand, the substrate diversity provided by the SKiD database can help map the boundaries of enzyme promiscuity, can drive the development of predictive methods to identify non-native enzyme-substrate interactions (Kroll et al., 2023; Wang et al., 2024) and explain the structural basis behind substrate selectivity (Thakur and Pandit, 2022). The database can also lead to development of novel machine learning and deep learning models for accurate prediction of enzyme kinetic parameters, which can be used as *in silico* filters for enzyme design and mutation prioritization (Li et al., 2022; Kroll et al., 2023; Yu et al., 2023; Gollub et al., 2024; Shen et al., 2024; Wang et al., 2024; Wang et al., 2024; Cai et al., 2025). Since most of the existing methods are sequence-based, structure-based enzyme kinetics prediction models can leverage the SKiD database for training and validation. Since the database also records environmental conditions (pH and temperature) under which the kinetics data were recorded, the database and models developed using the data can enable improvement of existing enzyme-constrained metabolic models (ecModels) in predicting phenotypes more accurately (Chen et al., 2024).

## Conclusions

The structure of a protein dictates its function. The specific three-dimensional structure enables enzymes to bind to specific substrates with high specificity and catalyze biochemical reactions. Enzyme activity is usually measured through two kinetic parameters, k_cat_ and K_m_. Although there exist databases with experimental k_cat_ and K_m_ values for thousands of enzyme-substrate pairs, there is limited data to establish the relationship between these kinetic parameters and their corresponding enzyme-substrate three-dimensional structures. In this work, we attempted to bridge this gap through development of a high-quality structure-oriented kinetics database (SKiD) for enzyme-substrate pairs. SKiD is freely accessible through https://doi.org/10.5281/zenodo.15355031. The database includes 13,654 unique enzyme-substrate complexes, derived from 8,562 unique enzymes and 3,235 unique small molecular substrates, along with their k_cat_ and K_m_ values. The challenges faced during the database construction are also documented with examples, along with the measures taken to overcome those challenges. This resource will be potentially useful for predicting kinetic properties of novel engineered proteins for several different applications like green chemistry and metabolic engineering.

## Additional Information

The Supplementary Information 1 contains Tables S1-S3 and Figures S1-S10. The Supplementary Information 2 contains enzyme-PDB mapping, mutation mapping errors from BRENDA, outliers in the dataset and the dataset for docking benchmark.

## Author Contributions

**Sowmya Ramaswamy Krishnan**: Data Curation, Validation, Formal analysis, Software, Writing – Original Draft, Writing – Review & Editing, Visualization

**Nishtha Pandey**: Data Curation, Validation, Formal analysis, Software, Writing – Original Draft, Writing – Review & Editing

**Rajgopal Srinivasan**: Methodology, Validation, Writing – Review & Editing, Supervision

**Arijit Roy**: Conceptualization, Methodology, Validation, Investigation, Resources, Writing – Review & Editing, Supervision

## Competing Interests

All authors are employed by Tata Consultancy Services Limited.

## Acknowledgements

The authors sincerely thank Dr. Gopalakrishnan Bulusu, Dr. Navneet Bung, Mr. Dibyajyoti Das, Dr. Broto Chakrabarty, Mr. Sarveswara Rao Vangala, Mrs. Akriti Jain, Ms. Padmasini Raghavachary and Mr. Hrithik Gupta for their valuable suggestions and comments for the work.

## Notes

### Summary of Updates

Changed the author order. Arijit Roy, who is the corresponding author, will now appear as last instead of first.

