## Supplementary File 1 for "SKiD: A Structure-Oriented Kinetics Database of Enzyme-Substrate Interactions"

TCS Research (Life Sciences division), Tata Consultancy Services, Hyderabad 500081, India

^#^Authors with equal contribution

**Supplementary Information**

**Section S1: Effect of mutations on the enzyme kinetic parameters**

To understand the effect of mutations on K_m_ and k_cat_ values, 4,008 pairs of enzyme-substrate complexes were curated from SKiD with the condition that the activity of both the wild-type and mutant enzymes must have been tested against the same substrate (Supplementary Information 2). Only single-residue mutants were considered for this analysis, since the individual contribution of each residue cannot be directly delineated from multi-site mutation data. The mutations were categorized into one of 25 types, depending on the nature of the wild-type and mutant residues (aliphatic, polar uncharged, positively charged, negatively charged, aromatic). Among the binding site mutations, polar uncharged => aliphatic mutations were most prevalent while, negatively charged => aromatic mutations were least prevalent (Fig. S4a). Similarly, among the non-binding site mutations, polar uncharged => aliphatic mutations were most prevalent and aromatic => negatively charged mutations were least prevalent (Fig. S4b). In terms of the change in kinetic parameters post mutation, binding site mutations were found to induce higher average fold change in k_cat_ (Fig. S4c) and K_m_ (Fig. S4d) values compared to non-binding site mutations. In general, for both binding and non-binding site mutations, higher number of datapoints have positive pk_cat_ and negative pK_m_ difference. This could be a consequence of an intrinsic bias in the reporting of experimental observations, where enzyme engineering attempts which improve activity are considered as a success. An earlier study also raises similar concerns (Yan et al., 2022). The complete substitution matrix of binding and non-binding site mutations in the SKiD database is provided in the Supplementary Tables S1 and S2.

**Supplementary Tables**

**Table S1**: Substitution matrix with the percentage of all possible binding site mutations observed in the SKiD database.

| **WT/Mut** | **G** | **A** | **V** | **L** | **I** | **F** | **W** | **Y** | **M** | **S** | **C** | **T** | **P** | **N** | **Q** | **D** | **E** | **R** | **K** | **H** |
| --- | --- | --- | --- | --- | --- | --- | --- | --- | --- | --- | --- | --- | --- | --- | --- | --- | --- | --- | --- | --- |
| **G** | 0 | 1.53 | 0.08 | 0 | 0 | 0.04 | 0 | 0 | 0 | 0.22 | 0.19 | 0 | 0 | 0.37 | 0 | 0.22 | 0.11 | 0.63 | 0 | 0 |
| **A** | 0.9 | 0 | 0.22 | 0.15 | 0.34 | 0.11 | 0 | 0.19 | 0 | 0.37 | 0.52 | 0.19 | 0.04 | 0 | 0 | 0 | 0 | 0.52 | 0 | 0.15 |
| **V** | 0.19 | 0.97 | 0 | 0.45 | 0.26 | 0.37 | 0 | 0.22 | 0.04 | 0.08 | 0 | 0 | 0 | 0.11 | 0 | 0.04 | 0.04 | 0.08 | 0.15 | 0 |
| **L** | 0.26 | 0.86 | 0.52 | 0 | 0 | 0.26 | 0.08 | 0.22 | 0 | 0 | 0.04 | 0 | 0.19 | 0 | 0.04 | 0.04 | 0 | 0.04 | 0 | 0 |
| **I** | 0.04 | 0.6 | 0.75 | 0.08 | 0 | 0.11 | 0 | 0 | 0 | 0 | 0 | 0.22 | 0 | 0.15 | 0.08 | 0 | 0 | 0 | 0 | 0 |
| **F** | 0 | 1.64 | 0.37 | 0.75 | 0.22 | 0 | 1.01 | 0.63 | 0.11 | 0.26 | 0 | 0 | 0.08 | 0.3 | 0.08 | 0.04 | 0 | 0 | 0 | 0.04 |
| **W** | 0 | 1.38 | 0 | 1.23 | 0.15 | 0.9 | 0 | 0.41 | 0 | 0 | 0 | 0 | 0 | 0 | 0 | 0.04 | 0 | 0 | 0 | 0.19 |
| **Y** | 0.34 | 1.42 | 0.08 | 0.6 | 0.04 | 7.13 | 0.41 | 0 | 0.15 | 0.34 | 0.26 | 0.08 | 0 | 0.08 | 0 | 0.49 | 0.04 | 0.22 | 0 | 0.15 |
| **M** | 0.45 | 0.26 | 0.19 | 0.11 | 0.41 | 0.08 | 0 | 0 | 0 | 0.22 | 0.19 | 0.08 | 0 | 0 | 0.04 | 0.04 | 0.04 | 0.86 | 0 | 0.19 |
| **S** | 0.26 | 4.66 | 0.15 | 0 | 0.04 | 0 | 0 | 0 | 0 | 0 | 0.08 | 0.26 | 0.41 | 0.08 | 0 | 0.49 | 0.26 | 0 | 0 | 0 |
| **C** | 0.11 | 0.63 | 0.19 | 0.04 | 0 | 0 | 0 | 0 | 0 | 1.27 | 0 | 0 | 0.15 | 0 | 0 | 0.08 | 0 | 0.04 | 0 | 0 |
| **T** | 0.45 | 1.98 | 0.49 | 0.22 | 0.11 | 0.19 | 0 | 0.08 | 0 | 0.82 | 0.04 | 0 | 0 | 0.08 | 0.08 | 0.08 | 0.11 | 0 | 0 | 0 |
| **P** | 0 | 0.15 | 0 | 0 | 0 | 0 | 0 | 0 | 0 | 0 | 0.11 | 0.08 | 0 | 0 | 0 | 0 | 0 | 0 | 0 | 0 |
| **N** | 0 | 2.24 | 0.04 | 0.15 | 0 | 0.04 | 0 | 0.22 | 0 | 0.19 | 0.45 | 0.93 | 0 | 0 | 0.08 | 0.41 | 0.08 | 0.11 | 0 | 0.11 |
| **Q** | 0 | 0.56 | 0.08 | 0.11 | 0.08 | 0 | 0 | 0.08 | 0 | 0 | 0 | 0 | 0 | 0.63 | 0 | 0.11 | 1.38 | 0.63 | 0 | 0.15 |
| **D** | 0.26 | 4.37 | 0.08 | 0 | 0 | 0.04 | 0 | 0.04 | 0.04 | 0.9 | 0.04 | 0 | 0 | 3.55 | 0 | 0 | 1.12 | 0.04 | 0 | 0 |
| **E** | 0.19 | 2.43 | 0 | 0.45 | 0 | 0 | 0 | 0 | 0 | 0.04 | 0 | 0 | 0 | 0.11 | 1.6 | 0.86 | 0 | 0 | 0.49 | 0.37 |
| **R** | 1.64 | 4.03 | 0 | 0 | 0.22 | 0.08 | 0.04 | 0 | 0.37 | 0.3 | 0.71 | 0.22 | 0 | 0.04 | 0.49 | 0 | 0.22 | 0 | 1.75 | 0.41 |
| **K** | 0 | 2.76 | 0.22 | 0.49 | 0 | 0.08 | 0 | 0 | 0.11 | 0.04 | 0 | 0 | 0 | 0 | 0.45 | 0 | 0.15 | 0.56 | 0 | 0 |
| **H** | 0 | 2.58 | 0 | 0 | 0 | 0.11 | 0 | 0 | 0 | 0.37 | 0.11 | 0.19 | 0 | 1.12 | 0.71 | 0 | 0 | 0.04 | 0 | 0 |

**Table S2**: Substitution matrix with the percentage of all possible non-binding site mutations observed in the SKiD database.

| **WT/Mut** | **G** | **A** | **V** | **L** | **I** | **F** | **W** | **Y** | **M** | **S** | **C** | **T** | **P** | **N** | **Q** | **D** | **E** | **R** | **K** | **H** |
| --- | --- | --- | --- | --- | --- | --- | --- | --- | --- | --- | --- | --- | --- | --- | --- | --- | --- | --- | --- | --- |
| **G** | 0 | 1.05 | 0 | 0 | 0 | 0 | 0 | 0 | 0 | 0.3 | 0 | 0.08 | 0 | 0 | 0 | 0.68 | 0.23 | 0.45 | 0 | 0 |
| **A** | 0.45 | 0 | 0.08 | 0.6 | 0.08 | 0 | 0.15 | 0.68 | 0 | 0.08 | 0 | 0.38 | 0 | 0 | 0 | 0 | 0 | 0.15 | 0 | 0 |
| **V** | 0.23 | 0.23 | 0 | 0.3 | 0.3 | 0.08 | 0 | 0 | 0.75 | 0 | 0 | 0.23 | 0 | 0 | 0 | 0.15 | 0 | 0 | 0 | 0 |
| **L** | 0.3 | 1.21 | 0.23 | 0 | 0.3 | 0.23 | 0 | 0 | 0.15 | 0 | 0.08 | 0 | 0.08 | 0 | 0.3 | 0.45 | 0 | 0.3 | 0.08 | 0 |
| **I** | 0.08 | 0 | 0.75 | 0.15 | 0 | 0 | 0 | 0 | 0 | 0 | 0 | 0.3 | 0 | 0 | 0 | 0.45 | 0 | 0.15 | 0 | 0 |
| **F** | 0.08 | 1.51 | 0.3 | 1.28 | 0.08 | 0 | 0 | 1.21 | 0 | 0.08 | 0.45 | 0 | 0.83 | 0 | 0 | 0 | 0 | 0.23 | 0.45 | 0.08 |
| **W** | 0.08 | 0.75 | 0 | 0.75 | 0 | 1.51 | 0 | 0.08 | 0 | 1.43 | 0 | 0 | 0 | 0 | 0 | 0 | 0 | 0 | 0 | 0.83 |
| **Y** | 0 | 1.96 | 0 | 0.75 | 0 | 2.18 | 0.08 | 0 | 0.23 | 0.08 | 0.15 | 0.08 | 0 | 0 | 0 | 0 | 0 | 0 | 0 | 0.98 |
| **M** | 0 | 0.23 | 0 | 0 | 0.53 | 0.3 | 0 | 0 | 0 | 0 | 0 | 0 | 0 | 0 | 0 | 0 | 0 | 0 | 0 | 0.08 |
| **S** | 0.38 | 1.51 | 0 | 0.6 | 0 | 0.3 | 0 | 0 | 0 | 0 | 0.08 | 0.75 | 0.53 | 0.53 | 0.15 | 0.9 | 1.13 | 0 | 0 | 0.15 |
| **C** | 0 | 4.37 | 0 | 0 | 0 | 0 | 0 | 0.3 | 0 | 1.96 | 0 | 0 | 0 | 0 | 0 | 0.15 | 0 | 0 | 0 | 0 |
| **T** | 0 | 0.98 | 0.3 | 0.08 | 0.38 | 0 | 0.68 | 0 | 0 | 0.3 | 0 | 0 | 0 | 0.23 | 0 | 0.38 | 0.15 | 0.08 | 0 | 0 |
| **P** | 0.98 | 0.75 | 0 | 0.9 | 0 | 0 | 0 | 0 | 0 | 0.68 | 0 | 0.23 | 0 | 0 | 0.08 | 0 | 0 | 0 | 0 | 0.08 |
| **N** | 1.05 | 1.36 | 0.15 | 0 | 0 | 0 | 0.98 | 0.08 | 0 | 0.6 | 0 | 0.45 | 0 | 0 | 1.88 | 0.45 | 0.08 | 0.08 | 0.3 | 0 |
| **Q** | 0 | 0.53 | 0.08 | 0.3 | 0 | 0.08 | 0 | 0 | 0 | 0.15 | 0 | 0 | 0 | 0.15 | 0 | 0 | 0.15 | 0.23 | 0.98 | 0.23 |
| **D** | 0.83 | 3.31 | 0.15 | 0.08 | 0 | 0 | 0 | 0 | 0.23 | 0.15 | 0.53 | 0 | 0 | 2.49 | 0.3 | 0 | 0.08 | 0.68 | 0 | 0.45 |
| **E** | 0.45 | 1.96 | 0.08 | 0.08 | 0 | 0.23 | 0.08 | 0 | 0 | 0.3 | 0.15 | 0 | 0 | 0 | 0.9 | 0.83 | 0 | 0.08 | 1.13 | 0 |
| **R** | 0.38 | 1.88 | 0.15 | 1.21 | 0.15 | 0 | 0.75 | 0 | 0.08 | 0.15 | 0.08 | 0 | 0 | 0 | 0.68 | 0 | 0.38 | 0 | 1.21 | 0.75 |
| **K** | 0.08 | 1.13 | 0 | 0 | 0 | 0 | 0 | 0 | 0.15 | 0.15 | 0.38 | 0 | 0 | 0.53 | 1.88 | 0 | 0.53 | 0.3 | 0 | 0 |
| **H** | 0 | 3.99 | 0 | 0.08 | 0 | 0.08 | 0.15 | 0.38 | 0 | 0.15 | 0 | 0 | 0 | 1.51 | 0.53 | 0.23 | 0.23 | 0.08 | 0 | 0 |

**Table S3**: Comparative results summarizing all atom RMSD value (Å) of mutant structures predicted using different computational methods with respect to their experimental structure. The RMSD values were computed using the ProFit server.

| **Query (WT)** | **Mutation** | **All atoms RMSD (Å) of predicted versus experimental structure** | | | **Experimental structure** |
| --- | --- | --- | --- | --- | --- |
|  |  | **FASPR (sidechain repacking of mutated residue only)** | **FASPR (sidechain repacking of all residues)** | **SCWRL** |  |
| 1R37 | N249Y | 2.927 | 3.044 | 2.94 | 1NTO |
| 1RJW | L176F | 0.93 | 1.177 | 0.941 | 3PII |
| 1MI3 | H114A | 0.516 | 0.943 | 0.58 | 1R38 |
| 2XSR | L69A | 0.333 | 1.122 | 0.459 | 2XSV |
| 1H82 | K300M | 0.649 | 0.986 | 0.731 | 3KPF |
| 1DP0 | H418N | 0.497 | 1.143 | 0.638 | 3DYO |
| 1AMP | S228A | 0.643 | 0.988 | 0.672 | 3B3V |
| 3B3X | E166C | 2.127 | 2.21 | 2.135 | 3SH7 |
| 6SIR | E497K | 1.223 | 1.555 | 1.267 | 5OYH |
| 4C9S | H33E | 0.765 | 1.127 | 0.829 | 8B7U |

**Supplementary Figures**

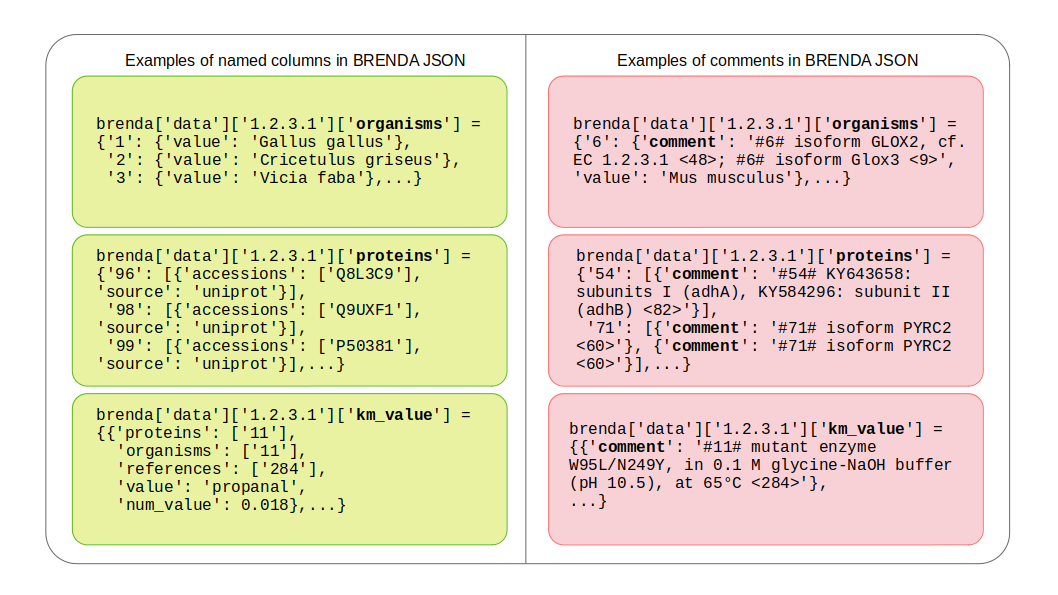
**Figure S1**: Examples of named columns and comments from BRENDA JSON file for the enzyme with EC number 1.2.3.1. A few entries under organisms, proteins and km_value categories in the JSON file are highlighted.

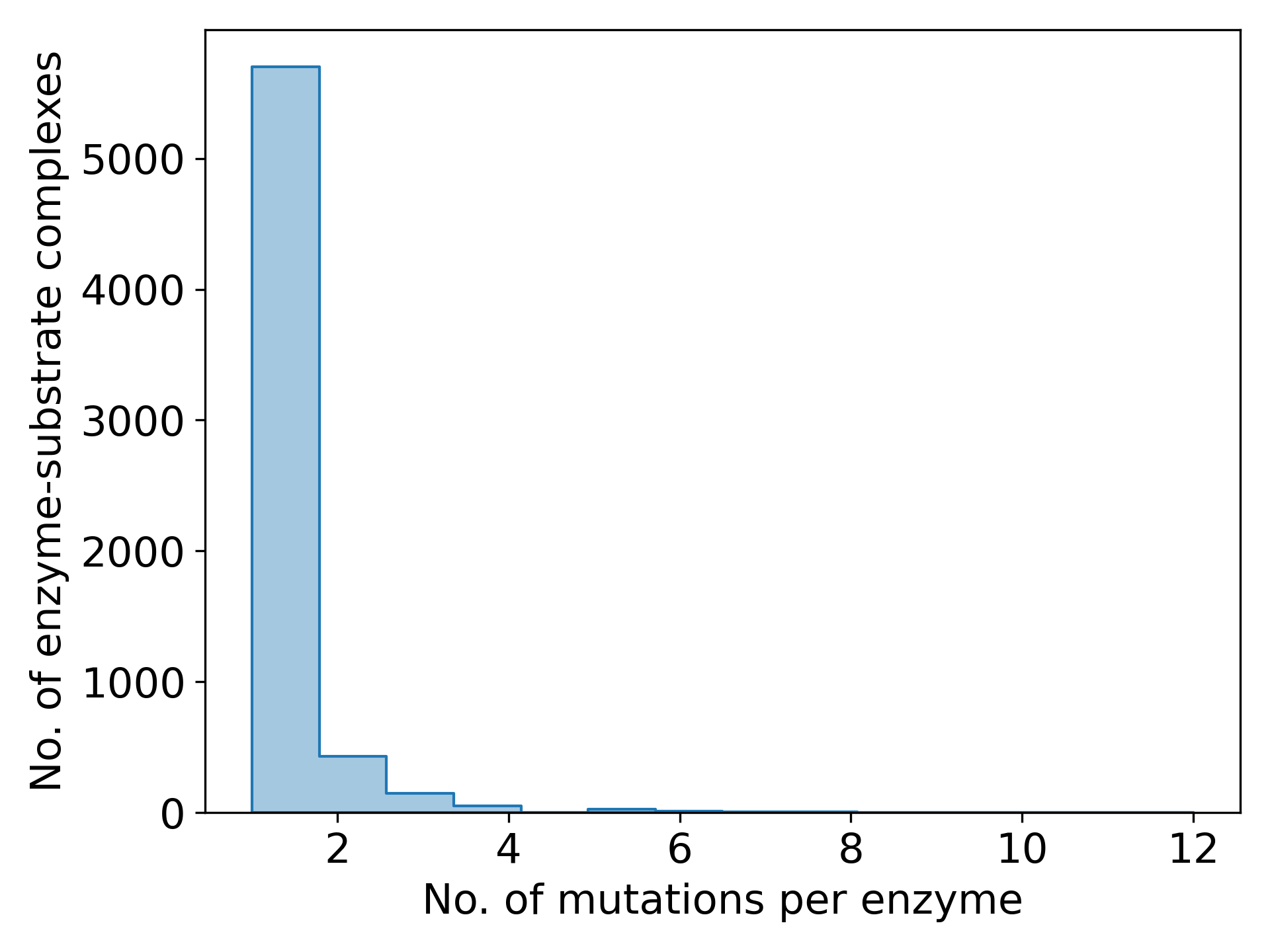
**Figure S2**: Histogram of the number of mutations screened simultaneously per enzyme in the BRENDA database.

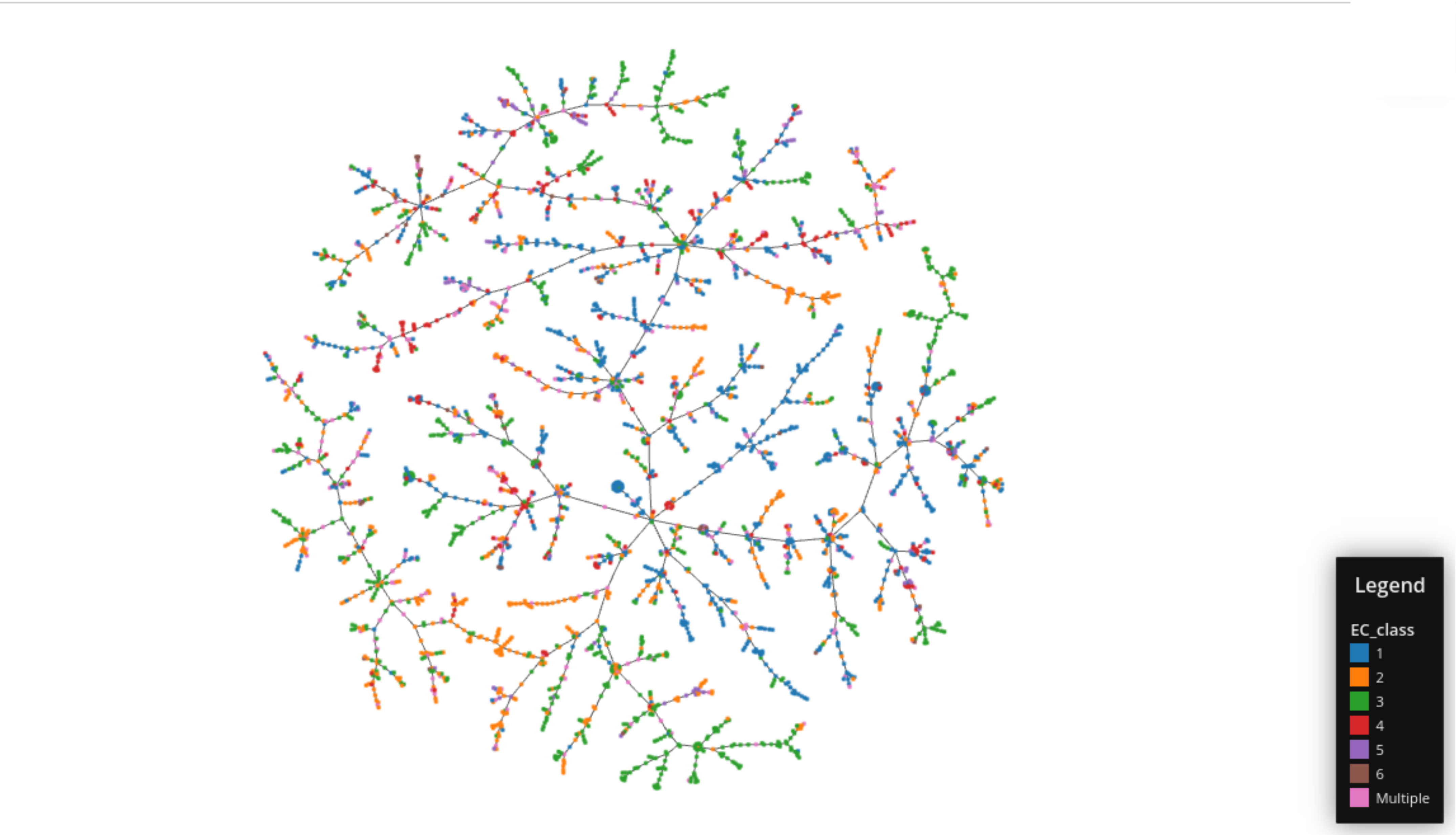
**Figure S3**: TMAP of the unique substrates present in BRENDA database colored based on the combination of EC classes with which they were observed to occur.

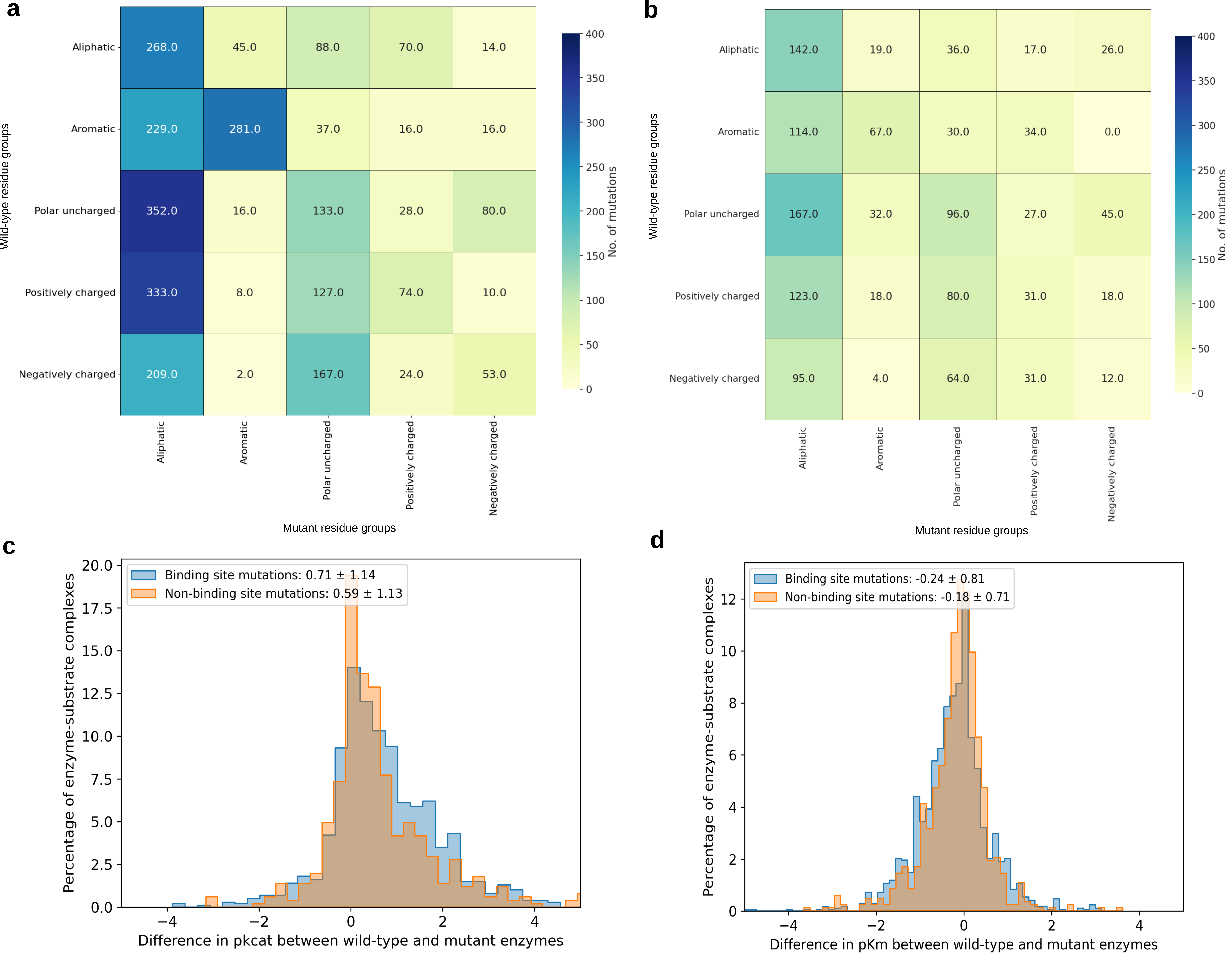
**Figure S4**: Heatmaps depicting the prevalence of 25 mutation types across single-residue mutants in the SKiD database among (a) binding site mutations and (b) non-binding site mutations. Distributions comparing the change in (c) pk_cat_ (=) and (d) pK_m_ (=) values between wild-type and mutant enzymes due to binding site and non-binding site mutations.

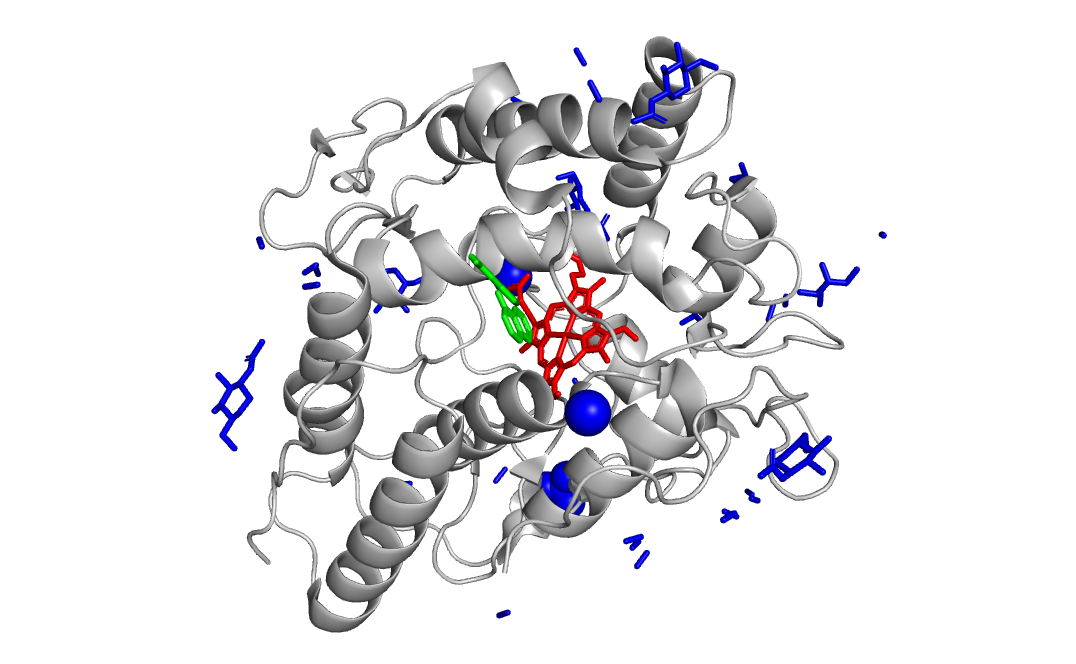
**Figure S5**: Example of an enzyme with multiple bound ligands – Unspecific peroxygenase (EC 1.11.2.1) from *Cyclocybe aegerita*. The substrate 1-naphthol is shown in green and the cofactor Heme is shown in red. All other ligands present in the structure including ions, carbohydrates and crystallization solvents are shown in blue. Except the substrate and cofactor, the other ligands were not considered in the database.

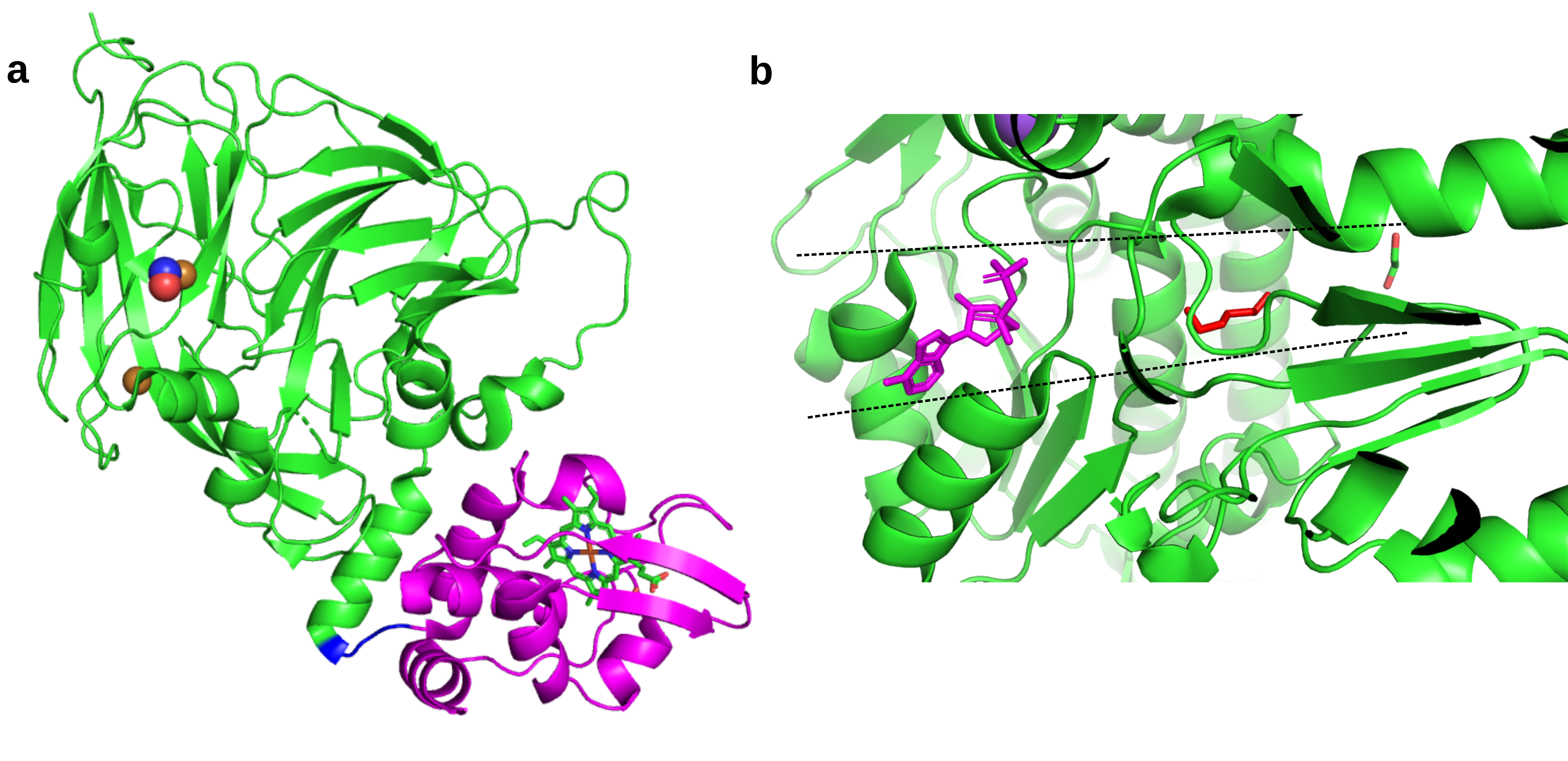
**Figure S6**: Distal substrate binding sites observed in cofactor-only PDB structures (**a**) Heme-Cu nitrite reductase (EC 1.7.2.1 – PDB ID: 5OCF) with distinct domains for cofactor (magenta) and substrate (green) binding; (**b**) Aminoaldehyde dehydrogenase (EC 1.2.1.19 – PDB ID: 4I9B) with a tunnel-shaped binding site (dashed lines) wherein the cofactor (magenta) and substrate (red) are bound to extreme ends of the tunnel.

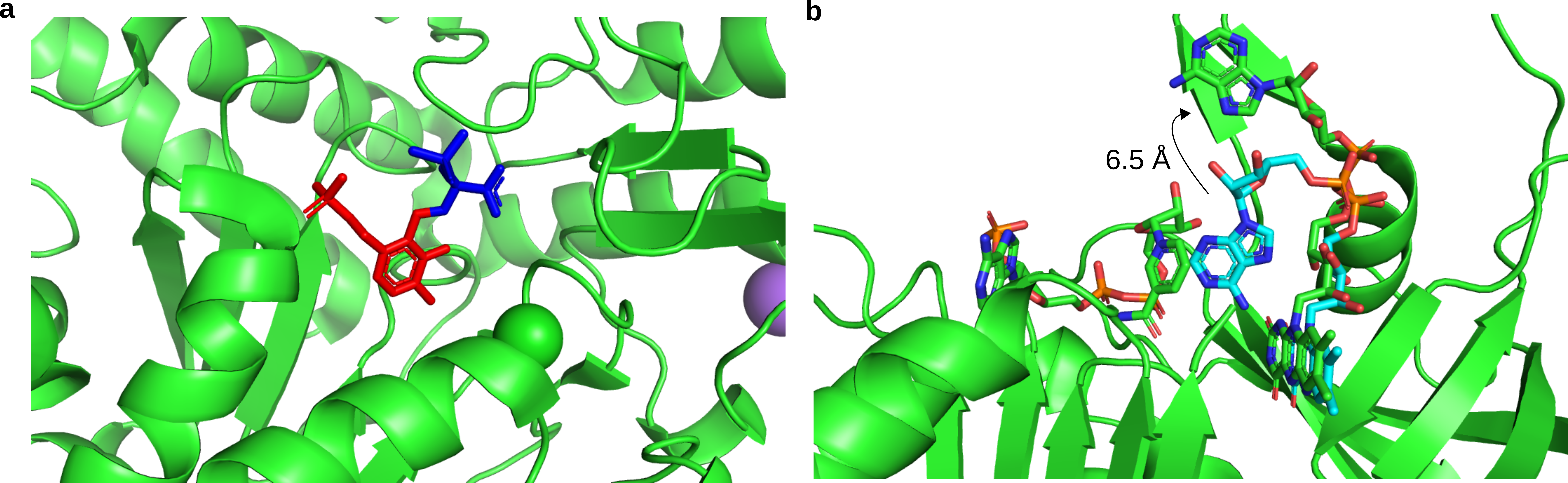
**Figure S7**: Proximal substrate binding sites observed in cofactor-only PDB structures (**a**) Phosphoserine aminotransferase (EC 2.6.1.52 – PDB ID: 4AZJ) with covalent C-N bond formation between the substrate (blue) and cofactor (red); (**b**) Ferridoxin-NADP+ reductase (EC 1.18.1.2) showing significant movement of the cofactor head group between the cofactor-only (PDB ID: 4B4D - green) and substrate+cofactor (PDB ID: 1GJR - cyan) forms of the enzyme.

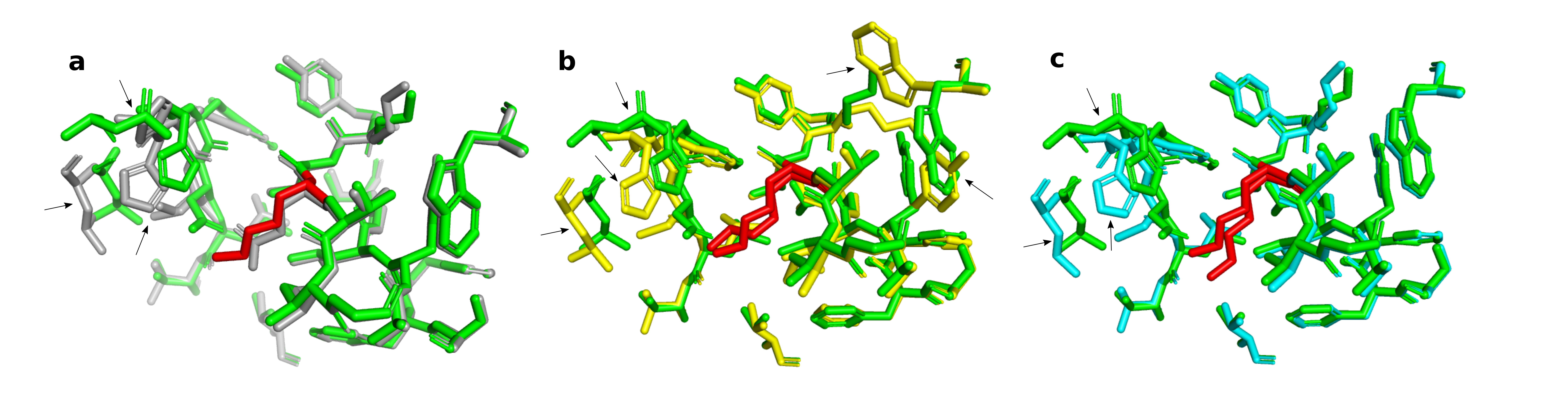
**Figure S8**: Effect of different residue repacking modes available in FASPR on the mutant structure generated from the wild-type enzyme – (**a**) Comparison between the experimental wild-type (PDB ID: 6SIR in gray) and mutant (PDB ID: 5OYH in green) structures with the active site residues showing difference in orientation highlighted with arrows due to E497K mutation (in red sticks); (**b**) Comparison between the experimental mutant (green) and mutant structure modelled using FASPR (yellow) with all position repacking option - residues Trp481, Lys495, Tyr503 (marked with arrows) show significant difference in side chain conformation; (**c**) Comparison between the experimental mutant (green) and mutant structure modelled using FASPR (cyan) with local residue repacking option.

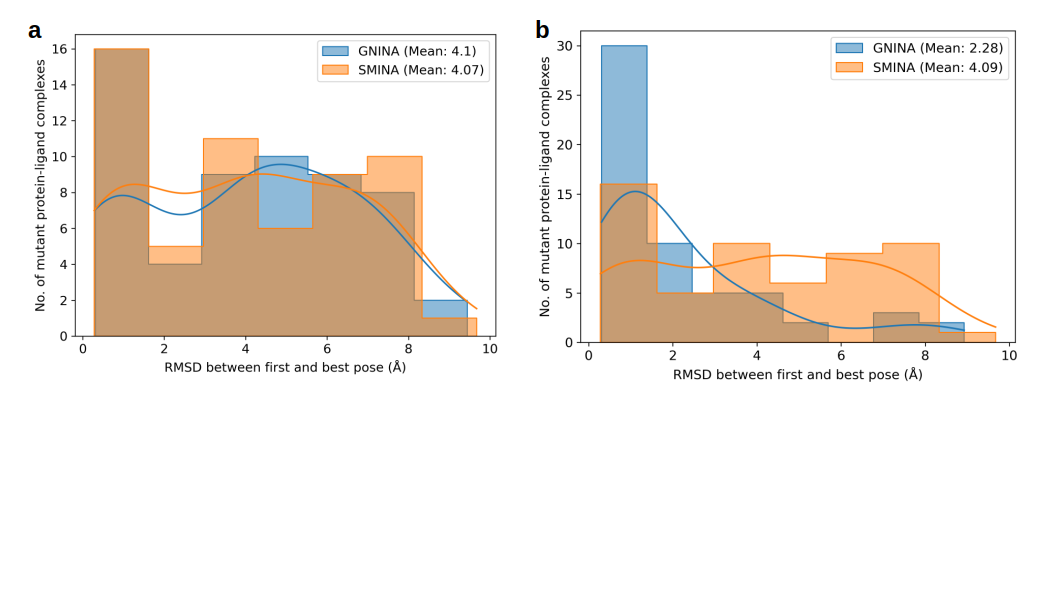
**Figure S9**: Comparison of the RMSD distributions (**a**) before rescoring and (**b**) after rescoring of the docking poses in GNINA for single pose sampling benchmark with the Platinum database. From the distributions, it is clear that CNN-based rescoring of docking poses significantly improves the RMSD distribution of GNINA compared with SMINA.

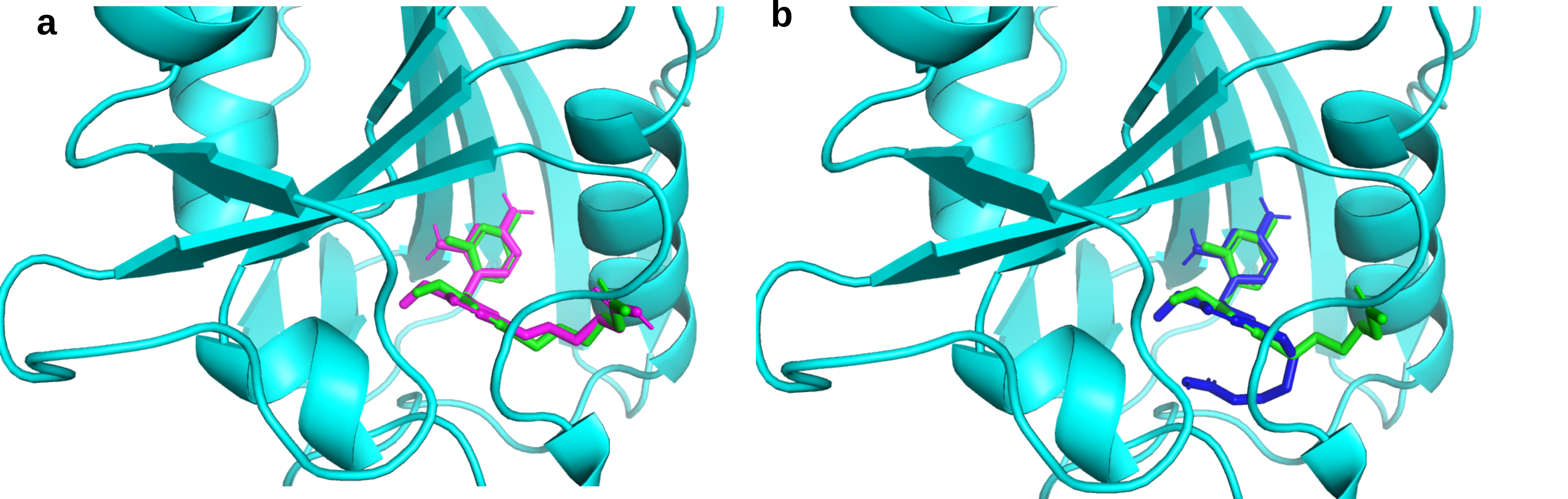
**Figure S10:** Docking of the ligand DH1 (green) to human dihydrofolate reductase (PDB ID: 3FS6) using (**a**) GNINA (magenta) program and (**b**) SMINA (blue) program. While the top-scoring pose from GNINA achieved as RMSD of 0.72 Å compared with the crystal pose, the pose from SMINA had a high RMSD of 3.96 Å.
